# Biophysical and biochemical evidence for the role of acetate kinases (AckAs) in an acetogenic pathway in pathogenic spirochetes

**DOI:** 10.1101/2024.10.12.617992

**Authors:** Ranjit K. Deka, Shih-Chia Tso, Wei Z. Liu, Chad A. Brautigam

## Abstract

Unraveling the metabolism of *Treponema pallidum* is a key component to understanding the pathogenesis of the human disease that it causes, syphilis. For decades, it was assumed that glucose was the sole carbon/energy source for this parasitic spirochete. But the lack of citric-acid-cycle enzymes suggested that alternative sources could be utilized, especially in microaerophilic host environments where glycolysis should not be robust. Recent bioinformatic, biophysical, and biochemical evidence supports the existence of an acetogenic energy-conservation pathway in *T. pallidum* and related treponemal species. In this hypothetical pathway, exogenous D-lactate can be utilized by the bacterium as an alternative energy source. Herein, we examined the final enzyme in this pathway, acetate kinase (named TP0476), which ostensibly catalyzes the generation of ATP from ADP and acetyl-phosphate. We found that TP0476 was able to carry out this reaction, but the protein was not suitable for biophysical and structural characterization. We thus performed additional studies on the homologous enzyme (75% amino-acid sequence identity) from the oral pathogen *Treponema vincentii*, TV0924. This protein also exhibited acetate kinase activity, and it was amenable to structural and biophysical studies. We established that the enzyme exists as a dimer in solution, and then determined its crystal structure at a resolution of 1.36 Å, showing that the protein has a similar fold to other known acetate kinases. Mutation of residues in the putative active site drastically altered its enzyme activity. A second crystal structure of TV0924 in the presence of AMP (at 1.3 Å resolution) provided insight into the binding of one of the enzyme’s substrates. On balance, this evidence strongly supported the roles of TP0476 and TV0924 as acetate kinases, buttressing the existence of the acetogenic pathway in pathogenic treponemes.

## Introduction

*Treponema* is a genus of spirochetal bacteria whose members colonize or parasitize diverse species, ranging from arthropods to mammals (1). Among the most studied of these bacteria are human pathogenic species known to cause syphilis (*T. pallidum pallidum*), bejel (*T. pallidum endemicum*), and yaws (*T. pallidum pertenue*) (1,2). A number of dental pathogens also belong to this genus, including *T. denticola*, *T. socranskii*, and *T. vincentii* (1). Thus, members of this genus represent significant threats to human health and well-being, with syphilis in particular posting alarming increases in incidence in the United States recent years (3). Particularly concerning in the U.S. is the rise (3) in congenital syphilis (CS), in which an infected mother passes the disease to her child perinatally; this can result in miscarriages, infant mortality, or a host of medical challenges for newborns.

Combatting CS and other diseases caused by treponemes would be aided by comprehensive understanding of their respective physiologies. However, these bacteria can be difficult to study in the laboratory. Indeed, only recently has *T. pallidum* been successfully continuously cultured (4), and this has been achieved only in co-culture with human epithelial cells. We have therefore explored the unique features of the treponemal lifestyle via a different route: a combination of structural biology, solution biophysics, and biochemistry. By determining the crystal structures of treponemal proteins and gleaning their functions, we have uncovered theretofore unknown aspects of treponemal biology, including a tripartite ATP-independent periplasmic (TRAP) nutrient transporter with a unique accessory protein (5,6), a first-in-class purine-nucleoside transporter (7), and a protein apparently involved in metal homeostasis (8,9).

Among the findings of our studies are several that relate to the metabolic lifestyle of *T. pallidum*. Specifically, our results led to the hypothesis that the bulk of the organism’s redox biochemistry appears to be dependent on flavin-based cofactors (10). Evidence for this conclusion includes our discoveries in *T. pallidum* of an ABC transporter specific for riboflavin (11) and a periplasmic enzyme (TpFtp) that transfers flavin moieties from FAD to recipient proteins (12–14). Additional factors lending credence to this “flavin-centric” hypothesis come from examination of *T. pallidum*’s genome: the bacterium is remarkable for its dearth of iron-based redox enzymes, it harbors a putative flavin-utilizing (RNF) complex for the formation of a chemiosmotic gradient, and there appears to be a flavin-salvage pathway (10,15). Central to this hypothesis is the existence of an acetogenic pathway in the organism’s cytoplasm (Fig. 1). This energy-conservation pathway capitalizes on the catabolism of exogenous D-lactate, culminating in the substrate-level phosphorylation of ADP to ATP and producing acetate as a by-product. To date, we have used a combination of structural, biophysical, and biochemical characterizations to confirm the activities of two of the enzymes in this pathway, namely the D-lactate dehydrogenase (TpD-LDH; (16)) and the phosphotransacetylase (TpPta; (17)).

**Figure 1.**
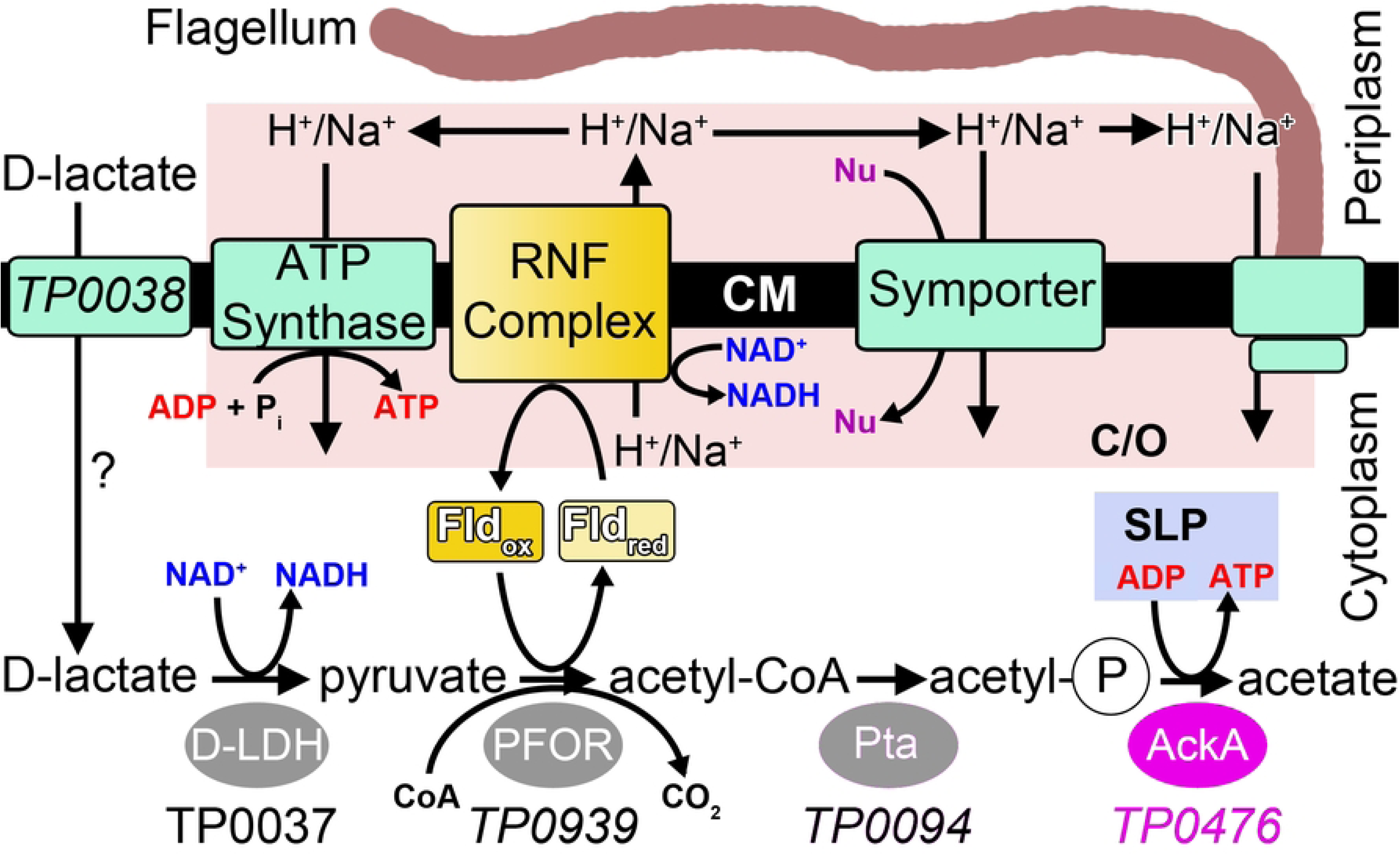
The hypothesized acetogenic pathway in *T. pallidum*. Relevant pathway enzymes are shown as ovals, with the enzyme that is featured in this work in purple. The equivalent of TP0476 in *T. vincentii* is TV0924 (not shown). Shaded yellow boxes depict flavoproteins, and cyan rectangles represent transmembrane proteins and complexes. The exception is the RNF complex, which is colored yellow to emphasize its status as a potential flavoprotein. The uncertain role of TP0038 in D-lactate import is noted with a question mark. The pink box labeled “C/O” encompasses proteins that ostensibly participate in the formation and utilization of the chemiosmotic gradient. The substrate-level phosphorylation putatively catalyzed by TP0476/TV0924 is in a light-blue box labeled “SLP.” Nu: nutrient; CM: cytoplasmic membrane. This figure was adapted (changes were made) from Fig 1 in (16). The license under which this was done is at https://creativecommons.org/licenses/by/4.0/.

In this report, we focus on the final enzyme in the pathway, i.e. the acetate kinase. In acetogenesis, the reaction catalyzed is:

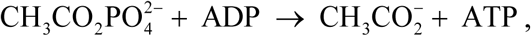

i.e., substrate-level phosphorylation of ADP using acetyl-phosphate as the phosphate donor. The *T. pallidum* protein that we identified as the putative acetate kinase is TP0476. In the process of overexpressing and purifying this protein, we discovered that its solution behavior was not ideal for the battery of experiments that we typically employ for a full characterization of an enzyme. We therefore chose to perform most of our studies on the comparatively well-behaved homolog from *T. vincentii* called “TV0924.” Our crystallographic, biochemical, and biophysical studies strongly support the notion that TV0924 and TP0476 are *bona fide* acetate kinases and hence likely participate in acetogenic pathways in their respective treponemes.

## Materials and Methods

### Materials

Unless otherwise noted below, all chemicals were purchased from Sigma-Aldrich (St. Louis, MO).

### Protein Expression, Purification, and Crystallization

To produce a recombinant derivative of TREVI0001_0924 (referred to in this section as “rTV0924”) in *Escherichia coli*, the DNA fragment encoding amino acid residues 1-449 of TV0924 was PCR amplified from *Treponema vincentii* ATCC700013 genomic DNA (ATCC, VA) by the polymerase incomplete primer extension (PIPE) cloning method using ends-specific primers (PIPE insert) (18). The expression vector, pSpeedET (DNASU, AZ), which encodes an N-terminal expression and purification hexa-histidine tag (MGSDKIHHHHHHENLYFQG), was PCR amplified with PIPE-vector primers. The PIPE-insert and PIPE-vector was mixed to anneal the amplified DNA fragments together (18). *E. coli* HK100 competent cells were transformed with the mixtures (PIPE-vector and insert) and selected for kanamycin resistance on LB agar plates. Cloning junctions/fragments were verified by DNA sequencing. A verified plasmid was then co-transformed with pGroESL (Takara) into *E. coli* BL21 AI (Invitrogen) cells for soluble protein expression. *E. coli* BL21 AI cells were grown at 37° C in LB medium containing 40 µg/mL of kanamycin and 30 µg/mL of chloramphenicol until the cell density reached an A600 of ∼0.6. The cells were then induced for ∼20 h with 0.2% (w/v) L-arabinose at 16 °C and harvested, and cell pellets were stored at −80 °C. The procedures for expression and purification of the recombinant proteins were essentially as previously described (5,13).

For the production of recombinant TP0476 (rTP0476) protein in *E. coli*, the DNA fragment encoding amino acid residues 1-448 of TP0476 was amplified from *Treponema pallidum* genomic DNA by the PIPE cloning method using ends-specific primers (PIPE insert) and cloned into a pSpeedET protein expression vector. The procedures for cloning, overproduction and purification of recombinant protein were essentially as described above.

For the construction of structure-guided rTV0924 variants, the N7A, R91A, H180A, R241A, and E388A, each mutation was individually introduced into the plasmid carrying the wild-type Tv0924 sequence using the QuikChange site-directed mutagenesis kit (Agilent Technologies). The mutation was confirmed by DNA sequencing. The mutant protein was expressed and purified as described above. Protein concentrations were determined in Buffer A (20 mM HEPES, 0.1 M NaCl, pH 7.5, 2 mM n-Octyl-β-D-glucopyranoside) using UV absorption at 280 nm. Extinction coefficients were calculated from the protein sequences using the ProtParam tool of ExPASy server (19).

Using the sitting-drop vapor-diffusion technique in 96-well plates, crystals of Apo-TV0924 were obtained after 3 months at 20 °C after the protein solution (at 60 mg/mL) was mixed 1:1 with the well solution, comprising 0.2 M (NH_4_)_2_SO_4_, 0.1 M Bis-Tris, pH 5.5, 25% (w/v) PEG 3350). Crystals of TV0924 were transferred to a stabilization solution comprising 0.2 M (NH_4_)_2_SO_4_, 0.1 M Bis-Tris pH 5.5, 25% PEG-3350, 0.1 M NaCl, 5% (v/v) ethylene glycol (EG), and 2 mM octyl-*β*-D-glucopyranoside. Subsequently, the crystals were transferred to the same solution with increasing concentrations of EG (15% and 25%) for approximately 2 min each, and then they were plunged into liquid nitrogen. They were stored at cryogenic temperatures until data collection. For AMP-TV0924, all of the conditions and procedures described above were identical except that the protein was at a concentration of 116 mg/mL and that 5 mM MgCl_2_ and 5 mM ADP were included at all crystallization, stabilization, and cryo-protection steps.

### Acetate Kinase Assays

The acetate kinase activity of TV0924 and TP0476 was assayed in the direction of ATP formation by measuring the generation of ATP from acetyl phosphate and ADP in the coupled assay with hexokinase and glucose-6-phosphate dehydrogenase (20). The 1-mL assay mixture contained 100 mM HEPES, pH 7.5, 100 mM NaCl, 20 mM MgCl_2_, 5.5 mM glucose, 1 mM NADP, 12 units of hexokinase, 6 units of glucose-6-phosphate dehydrogenase, and various concentrations of ADP and acetyl phosphate up to 5 mM and 10 mM, respectively. The assays were conducted in an Agilent 8453 diode-array UV Vis spectrophotometer by monitoring NADPH formation as increasing absorbance at 340 nm (ε_340_= 6.22 mM^−1^cm^−1^); the temperature-controlled cuvette holder was connected to a circulating water bath set to 37 °C. The enzymatic reaction was initiated by mixing TV0924 or TP0476 into the pre-incubated assay mixture. A 30 s stretch of a time trace between 50 and 80 s after the reaction initiated was fitted to a straight line and the slope was taken as the reaction rate. The reaction rates were maintained within 0.001-0.02 ΔA_340_/s by varying the enzyme amount (0.05-100 µg) added in any given assay. To estimate the *K*_m_ and *V*_max_, the acetate kinase activities at various substrate concentrations were fitted to the Michaelis-Menten equation using LabPlot2 (21). *V*_max_ (in units of µmol‧mg^−1^‧min^−1^ was converted to *k*_cat_ (in units of s^−1^) by the formula

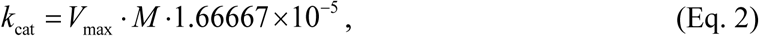

where *M* is molar mass of the respective protein. The assays were performed in triplicate (i.e., N = 3), and the values reported in this report are the weighted means of the respective parameters (*p̑*) according to the formula

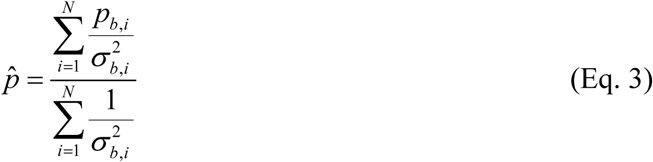

where *p*_b_ is the best value of the parameter fitted by LabPlot, and *σ_b_* is the respective error in *p* reported by LabPlot. The standard deviation of the mean for parameter *p̑*, *σ̑*, was calculated as

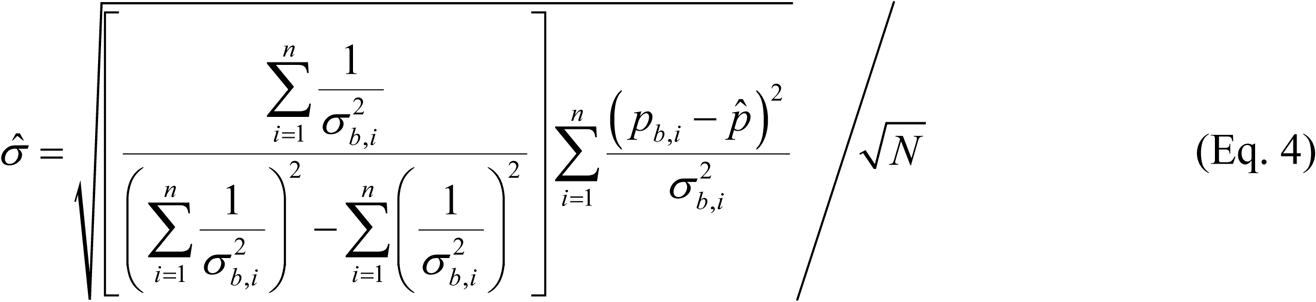

### Data Collection, Structure Determination, and Refinement

Cryo-cooled crystals of Apo-TV0924 and AMP-TV0924 were transported to synchrotron facilities for diffraction-data collection. Data sets from Apo-TV0924 crystals were acquired at 100 K at beamline 19-1 at the Structural Biology Center of the Advanced Photon Source at Argonne National Laboratories. The crystals had the symmetry of space group P2_1_2_1_2, and they diffracted X-rays to a *d*_min_ spacing of 1.36 Å (Table 1). To determine the structure, molecular replacement was employed, using a model generated by AlphaFold2 (22,23). The Phaser (24) implementation in Phenix (25) located a single molecule of TV0924 in the asymmetric unit with high confidence (Log-Likelihood Gain (LLG) = 2,045; Translation-Function Z (TFZ) = 37.9). The model was adjusted in Coot (26), and then the rigid-body refinement, simulated-annealing, positional, and *B*-factor refinement protocols in Phenix were utilized in an iterative fashion with model adjustment in Coot as necessary. Riding hydrogen atoms were used, and, in the late stages of refinement, anisotropic *B*-factors were refined. The final model had good fitting and geometric characteristics (Table 1).

**Table 1.**
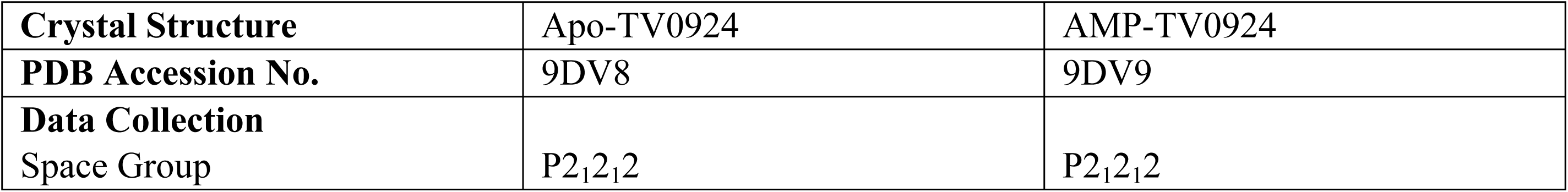
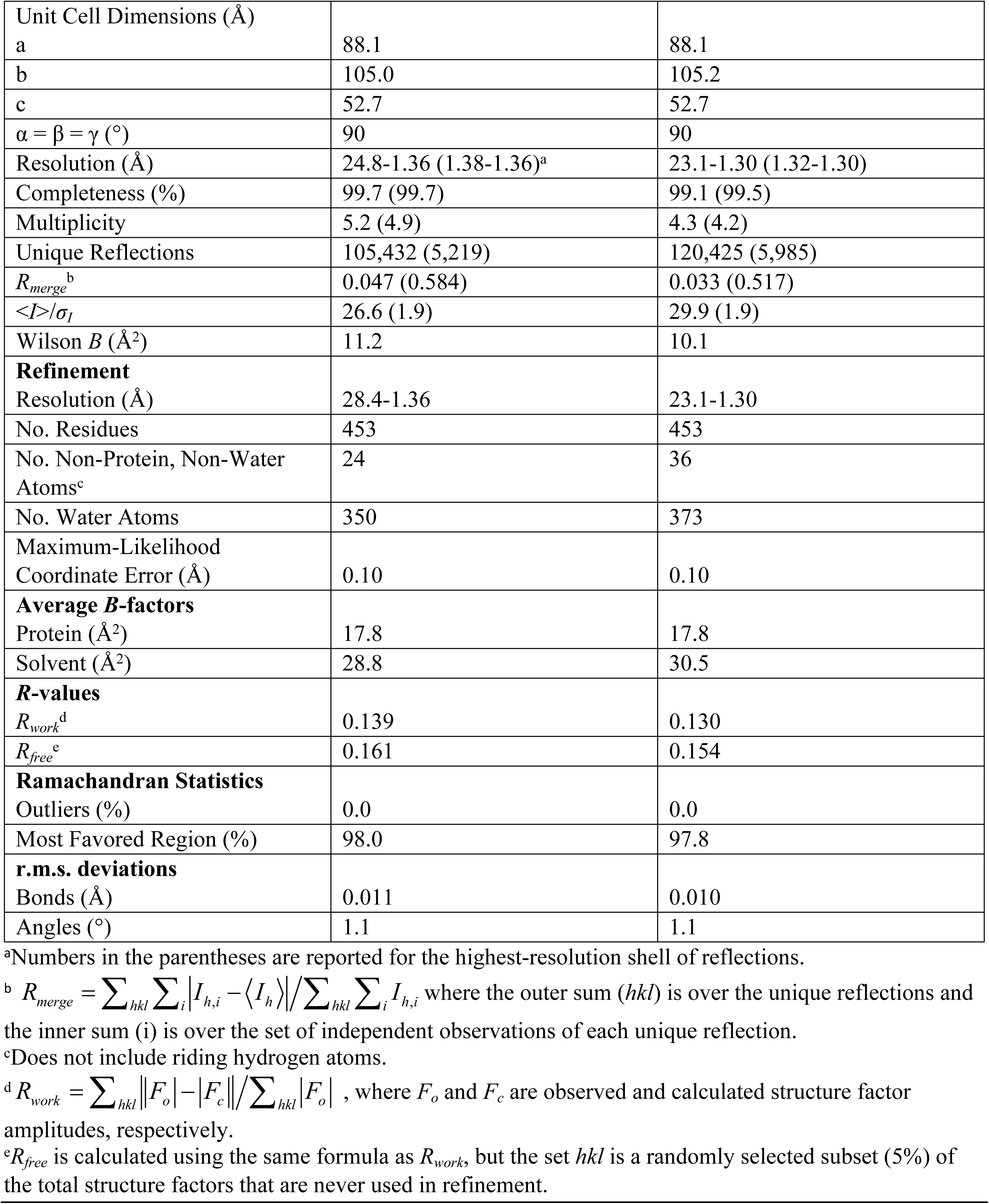
X-ray diffraction data and refinement statistics.

|  |  |  |
| --- | --- | --- |
| **Crystal Structure** | Apo-TV0924 | AMP-TV0924 |
| **PDB Accession No.** | 9DV8 | 9DV9 |
| **Data Collection** |  |  |
| Space Group | $P2_12_12$ | $P2_12_12$ |
| Unit Cell Dimensions (Å) |  |  |
| a | 88.1 | 88.1 |
| b | 105.0 | 105.2 |
| c | 52.7 | 52.7 |
| $\alpha = \beta = \gamma$ (°) | 90 | 90 |
| Resolution (Å) | 24.8-1.36 (1.38-1.36)<sup>a</sup> | 23.1-1.30 (1.32-1.30) |
| Completeness (%) | 99.7 (99.7) | 99.1 (99.5) |
| Multiplicity | 5.2 (4.9) | 4.3 (4.2) |
| Unique Reflections | 105,432 (5,219) | 120,425 (5,985) |
| $R_{merge}^b$ | 0.047 (0.584) | 0.033 (0.517) |
| $\langle I \rangle / \sigma_I$ | 26.6 (1.9) | 29.9 (1.9) |
| Wilson B (Å<sup>2</sup>) | 11.2 | 10.1 |
| **Refinement** |  |  |
| Resolution (Å) | 28.4-1.36 | 23.1-1.30 |
| No. Residues | 453 | 453 |
| No. Non-Protein, Non-Water Atoms<sup>c</sup> | 24 | 36 |
| No. Water Atoms | 350 | 373 |
| Maximum-Likelihood Coordinate Error (Å) | 0.10 | 0.10 |
| **Average B-factors** |  |  |
| Protein (Å<sup>2</sup>) | 17.8 | 17.8 |
| Solvent (Å<sup>2</sup>) | 28.8 | 30.5 |
| **R-values** |  |  |
| $R_{work}^d$ | 0.139 | 0.130 |
| $R_{free}^e$ | 0.161 | 0.154 |
| **Ramachandran Statistics** |  |  |
| Outliers (%) | 0.0 | 0.0 |
| Most Favored Region (%) | 98.0 | 97.8 |
| **r.m.s. deviations** |  |  |
| Bonds (Å) | 0.011 | 0.010 |
| Angles (°) | 1.1 | 1.1 |

<sup>a</sup>Numbers in the parentheses are reported for the highest-resolution shell of reflections.
<sup>b</sup> $R_{merge} = \sum_{hkl} \sum_i |I_{h,i} - \langle I_h \rangle| / \sum_{hkl} \sum_i I_{h,i}$ where the outer sum ( $hkl$ ) is over the unique reflections and the inner sum ( $i$ ) is over the set of independent observations of each unique reflection.
<sup>c</sup>Does not include riding hydrogen atoms.
<sup>d</sup> $R_{work} = \sum_{hkl} ||F_o| - |F_c|| / \sum_{hkl} |F_o|$ , where $F_o$ and $F_c$ are observed and calculated structure factor amplitudes, respectively.
<sup>e</sup> $R_{free}$ is calculated using the same formula as $R_{work}$ , but the set $hkl$ is a randomly selected subset (5%) of the total structure factors that are never used in refinement.

Data sets from crystals of AMP-TV0924 were collected at 100 K at beamline 12-2 at the Stanford Synchrotron Radiation Lightsource at the SLAC National Accelerator Laboratory. These crystals had the same symmetry as the apo-crystals, and they diffracted X-rays to a similar *d*_min_ spacing (1.30 Å). The same strategy as used for Apo-TV0924 for solving the phase problem was followed, with similarly robust initial results (LLG = 2,067, TFZ = 37.9). Initial difference electron-density maps were closely examined for the presence of Mg(II) and ADP, but only AMP could be located. The nucleotide was modeled using Coot, and refinement proceeded as outlined above for the Apo-TV0924 structure. As before, the model exhibited good fitting and geometry statistics (Table 1). A region on the periphery of the protein (residues 251-272) showed evidence for two discrete conformations, and it was therefore modeled as such.

### Analytical Ultracentrifugation

All analytical ultracentrifugation (AUC) experiments were conducted in the sedimentation velocity (SV) mode in a Proteome XL-I centrifuge (Beckman-Coulter, Indianapolis, IN). Centrifugation cell assemblies were prepared by sandwiching a standard, charcoal-filled Epon centerpiece between two sapphire windows in an aluminum housing. All experiments were performed at 20 °C in Buffer A. Approximately 400 µL of the sample was inserted into the sample sector of the centerpiece, and the same volume of Buffer A was introduced into the reference sector. After sealing the fill-port holes, the assemblies were positioned into an An-50Ti rotor, which was subsequently placed into the centrifuge. Chamber evacuation and temperature equilibration were then undertaken for approximately 2.5 h. Next, centrifugation at 50,000 rpm was initiated, and data acquired using the absorbance optical system tuned to 280 nm. Centrifugation continued until no sedimenting components could be observed. SV data were analyzed using SEDFIT’s *c*(*s*) distribution (27,28). Time-invariant noise decomposition (29) was employed, along with a resolution of 150, a maximum-entropy regularization level of 0.683, an *s*_min_ of 0.0 S, and an *s*_max_ of 25 S. The sample meniscus position and the frictional ratio were refined. The proteins’ partial-specific volumes, the solution density, and the solution viscosity were calculated using SEDNTERP (30,31). Integrations of the *c*(*s*) distributions from SEDFIT and rendering of SV figures were performed in GUSSI (32).

### Mass Photometry

Mass photometry (MP) experiments were conducted using a TwoMP apparatus (Refeyn, Oxford, UK). Automated focusing operations were performed after placing 16.2 µL of phosphate-buffered saline (PBS) solution into a silicone-gasket well on a glass cover slide that had been situated on the instrument. To this, 1.8 µL of sample (diluted to 100 nM in PBS) was added, and the droplet was mixed by pipetting. A 60-s movie of the interferometric signal from an area (ca. 46 µm^2^ was acquired. Using the Discover^MP^ software, this movie was converted to a ratiometric form, and contrast events were characterized and tabulated by the software. The software also related contrast values to molecular masses via a calibration curve that had been previously constructed by collected data on the monomers and multimers observable in a sample of bovine serum albumen. These results were inputted into a custom Python script that formed histograms from the data and fitted Gaussian functions to selected peaks (33,34). The mean (µ) values from the Gaussian curves were taken as the molecular masses, and the *σ* values as their respective errors. Molecular masses reported herein were the weighted means of these results, and the errors in the means were calculated as the weighted standard deviation of the mean. These latter calculations were performed using (Eq. 3) and (Eq. 4).

### Circular Dichroism Spectrometry

Stability tests of the proteins were undertaken in a J-815 circular dichroism (CD) spectrometer (Jasco Inc., Easton, MD). Samples in Buffer A were diluted to 0.4 mg/mL, and then they were placed in a 1-mm quartz cuvette. After spectral characterization, the CD signal was monitored at 219 nm. The temperature of the sample was varied from 25 °C to 95 °C in increments of 1 °C by means of a Peltier temperature-control device (Jasco model CDF-426S). Data were collected using a slit width of 1 nm and a data-integration time of 4 s. Using a custom Python script, the resulting curves of CD vs. temperature were fitted to the equation

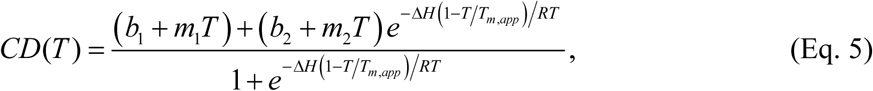

where *b*_1_ and *b*_2_ are the *y*-intercepts of the low-temperature and high-temperature parts of the curve, *m*_1_ and *m*_2_ are the respective slopes, Δ*H* is the molar enthalpy of unfolding, *T*_m,app_ is the apparent melting temperature, *T* is the temperature in Kelvins, and *R* is the gas constant. Apparent *T*_m_ values are reported because the reverse “melt” was not performed. Confidence intervals on the *T*_m,app_ values were procured using a confidence-interval search procedure built into the Python module lmfit.

## Results

### Purification, enzyme activities, and solution behavior of TP0476 and TV0924

Both TP0476 and TV0924 were cloned into a pSpeedET plasmid that positions a His_6_ affinity-purification tag and a tobacco etch virus (TEV) protease site at the amino-terminus of the recombinant protein. The recombinant proteins, which share 75% amino-acid identity (Fig. 2), expressed well in *E. coli*. Using a standard two-step preparation (affinity and size-exclusion chromatographies, see *Materials & Methods*), the proteins could be purified to near homogeneity. However, during subsequent steps to concentrate the proteins for storage, TP0476 could only be concentrated to about 10 mg/mL before it precipitated. On the other hand, TV0924 could be concentrated to about 60 mg/mL. We therefore focused most of our biochemical and biophysical studies on the latter enzyme.

**Figure 2.**
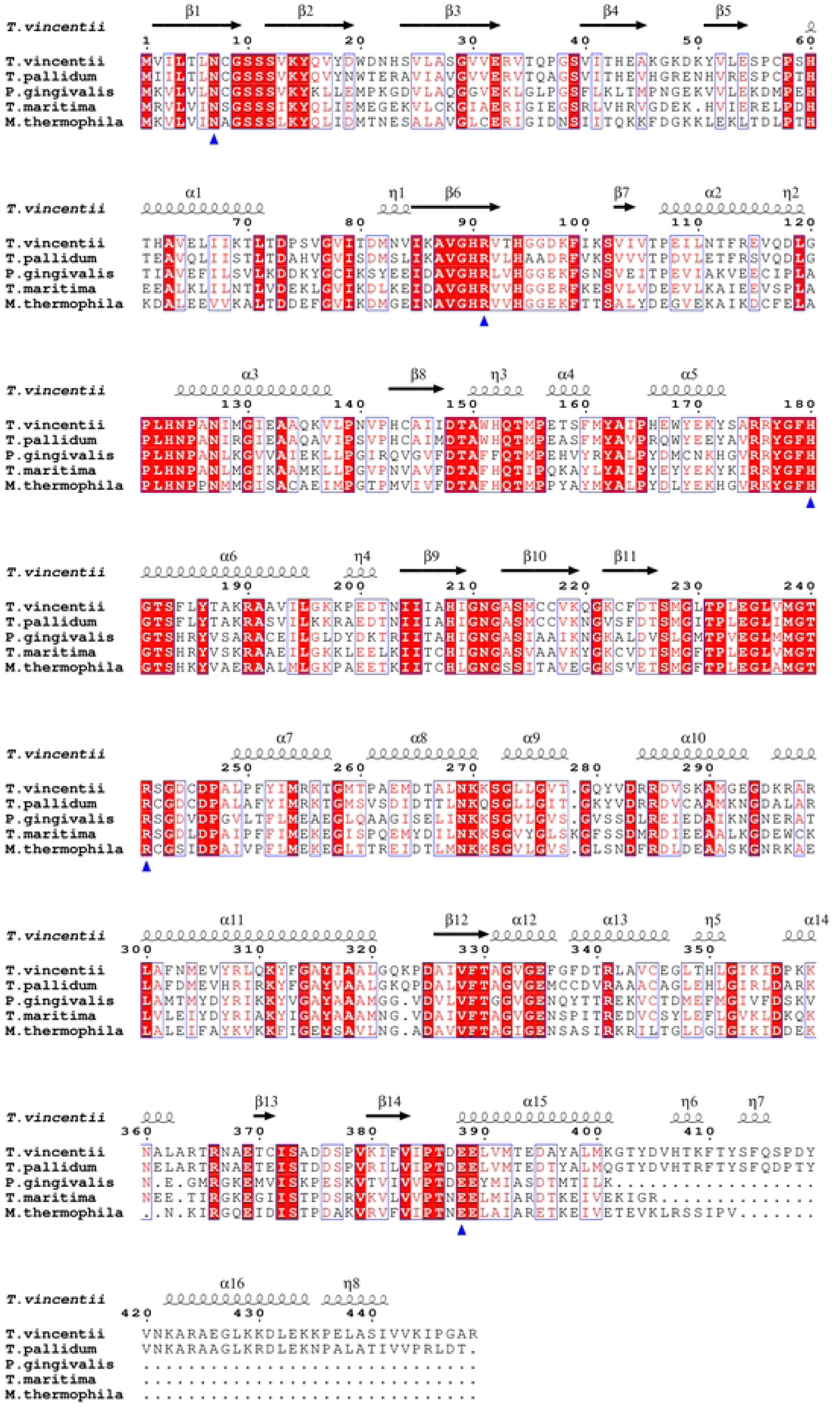
Sequence alignment of TV0924, TP0476, and acetate kinases. Shown are the single-letter amino acid codes for the *T. pallidum* and *T. vincentii* proteins, as well as the two best structural matches to TV0924 in the database as of this writing: AckA proteins from *P. gingivalis* and *T. maritima*. The secondary structures as assigned from the TV0924 structure is shown curled lines (α-or 3_10_-helices) and arrows (β-strands). Residues with a red background are absolutely conserved, while boxed residues have high similarity among the four proteins. Amino-acid residues marked with blue triangles are putative active-site residues that were mutated to alanine in this study, and the numbering shown is for the *T. vincentii* protein. This figure was generated using ESPRIPT version 3 (35).

To establish that the proteins were *bona fide* acetate kinases, i.e. that they are capable of substrate-level phosphorylation and acetate production (Fig. 1), we performed enzyme assays (Table 2). Acetyl phosphate and ADP were supplied, and the production of ATP was monitored (see *Materials & Methods*). Both proteins effectively catalyzed the reaction (Table 2) under our assay conditions, but the activity of TP0476 was substantially lower than its counterpart from *T. vincentii*; for example, the catalytic efficiency (*k*_cat_/*K*_M_) with respect to ADP was 7,500 mM^−1^s^−1^ for TV0924 vs. 90 mM^−1^s^−1^ for TP0476, a 83-fold difference. Remarkably, this difference is due solely to differences in *k*_cat_, as the *K*_M_ values are identical (Table 2).

**Table 2.**
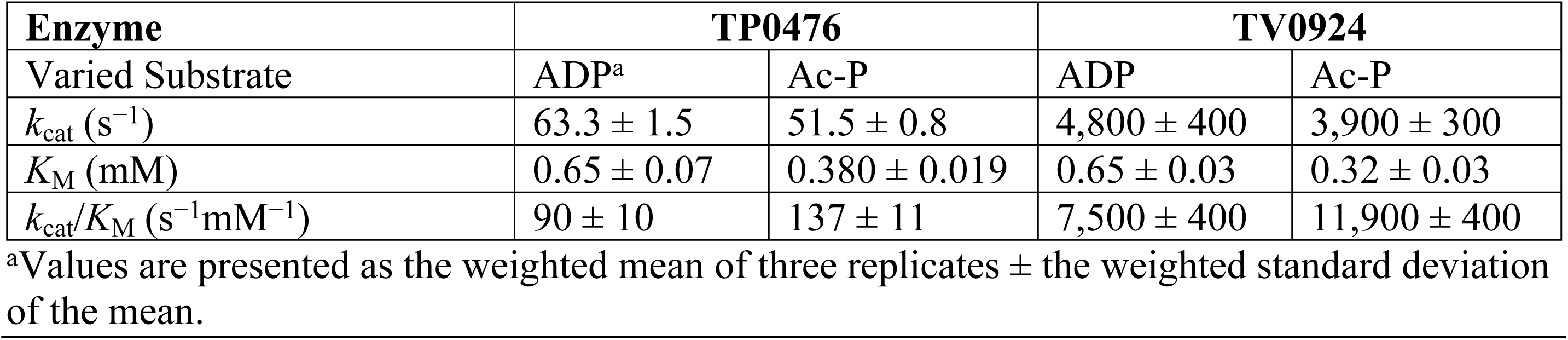
Enzymatic kinetic parameters for TP0476 and TV0924.

We hypothesized that the poorer solubility and activity of TP0476 compared to TV0924 would manifest as a difference when comparing the proteins’ respective solution behaviors and stabilities. We first studied the temperature stability of the proteins by monitoring their circular dichroism at a wavelength of 219 nm (Fig. 3A). TP0476 was clearly less stable than TV0924 at elevated temperatures, evincing a *T_m,_*_app_ that was about 14 °C lower.

**Figure 3.**
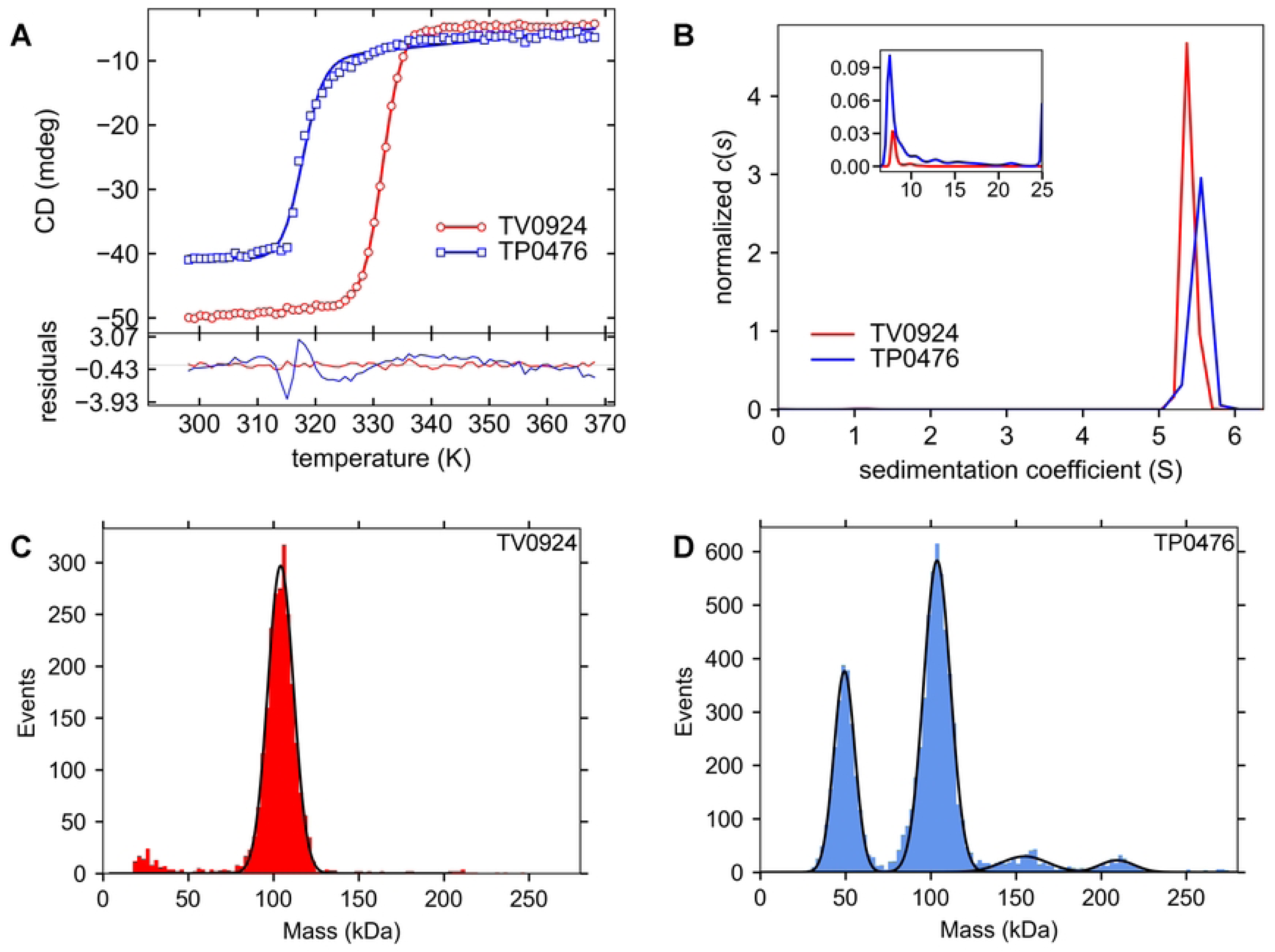
Biophysical characteristics of the TP0476 and TV0924. (A) Thermal stabilities of the proteins. In the *upper graph*, markers represent the circular dichroism values, and lines are fits to those values as described in Materials & Methods. Residuals between the data and the fit lines are shown in the *lower graph*. (B) Sedimentation velocity studies. Respective *c*(*s*) distributions are shown. The inset shows the higher *s*-range used for the analysis from about 6.5 S to 25 S. (C) A representative mass-photometry experiment for TP0476. The blue histogram represents the number of counts attributed to each given molecular mass. Black lines represent gaussian fits to the four peaks present in the histogram data. (D) A representative mass-photometry experiment for TV0924. Histogram and line conventions established in part (C) are followed, except the histogram is red. Low-molecular-mass readings were not fitted, as they are likely artifactual.

Next, using the data from the sedimentation velocity mode of analytical ultracentrifugation, we found that TP0476 displayed a dominant species at 5.49 ± 0.02 S (average ± standard deviation of the mean, N = 3; consistent with a dimer of the protein), but there was significant evidence of larger, aggregated forms (Fig. 3B) accounting for approximately 15% of the sedimenting signal. The molar mass for this protein was not calculated due to the unusually inconsistent frictional ratios refined among the replicates (1.19-1.29), which was likely due to the prevalence of the contaminants. On the other hand, TV0924 had almost no aggregates (2% of the signal), and its dominant species displayed a sedimentation coefficient of 5.387 ± 0.014 S with consistent frictional ratios (ranging from 1.32 to 1.36). Calculation (using the Svedberg equation) of the molar mass (*M*) with these values yielded *M* = 99,500 ± 600 g/mol *versus* a calculated molar mass of the dimer of 104,820 g/mol. It therefore appeared that the solution oligomeric states of the proteins were dimers, although TV0924 once again displayed evidence of enhanced stability.

As a final check on the *in vitro* behaviors of the proteins, we performed mass photometry (MP) (Figs. 3C & 3D). The resulting mass distribution for TP0476 (Fig. 3C) showed multiple peaks, with the main peak occurring at 103.43 ± 0.14 kDa. A species with a smaller mass (49.14 ± 0.11 kDa) was likely the monomeric form of the protein, while larger forms at 155.2 ± 0.3 kDa and 208.8 ± 0.3 kDa could be attributed to trimeric and tetrameric forms, respectively. Conversely, only a single peak could be observed consistently in MP data from TV0924 (Fig. 3D). The molecular masses derived from Gaussian fits to these peaks (104.58 ± 0.14 kDa) were highly reproducible and very close to the calculated value for a dimer (*vide supra*). It is striking that, even at the low concentration employed for MP (ca. 10 nM), the TV0924 dimer exhibits no tendency to dissociate into its protomers. To test this further, we incubated the protein for 65 h at 4 °C at 10, 20, and 30 nM, then performed MP (at ambient temperature; see Supplemental Fig. S1). Again, there was no evidence of monomer formation, indicating an extremely slow off-rate and likely a very low *K*_D_ for dimer formation in TV0924.

### The crystal structures of TV0924

We were not able to obtain crystals of recombinant TP0476, likely due to its poor stability and solution behavior (see above). However, TV0924 crystallized (albeit after a 90-day incubation period), and the crystals diffracted X-rays to a *d*_min_ spacing of 1.36 Å (Table 1). The structure was determined using molecular-replacement techniques employing a model generated by AlphaFold2 (22,23) from the amino-acid sequence of TV0924. A single copy of TV0924 was located in the asymmetric unit of the crystals, and the model, after refinement, exhibited excellent geometry (Table 1).

Overall, TV0924 adopts an α/β fold with two prominent β-sheets forming the respective centers of two lobes (Fig. 4). In keeping with conventions established with previously characterized acetate kinases, the lobes are called Domain I (the domain with the N-terminus, residues 1-148, 388-400) and Domain II (residues 149-387). A large, solvent-accessible cleft is located between the lobes, and previous studies on acetate kinases (36–39) and other sugar kinases (40) have identified this cleft as the enzyme’s active site. Indeed, residues thought to be critical for catalysis are located in the cleft in these other enzymes as well as TV0924. Intriguingly, the difference electron-density maps in the cleft region reveal evidence for a small molecule bound there (Fig. S2). However, the shape of this density did not match any probable substrate nor any molecule present in the crystallization medium. We chose to leave this density unmodeled.

**Figure 4.**
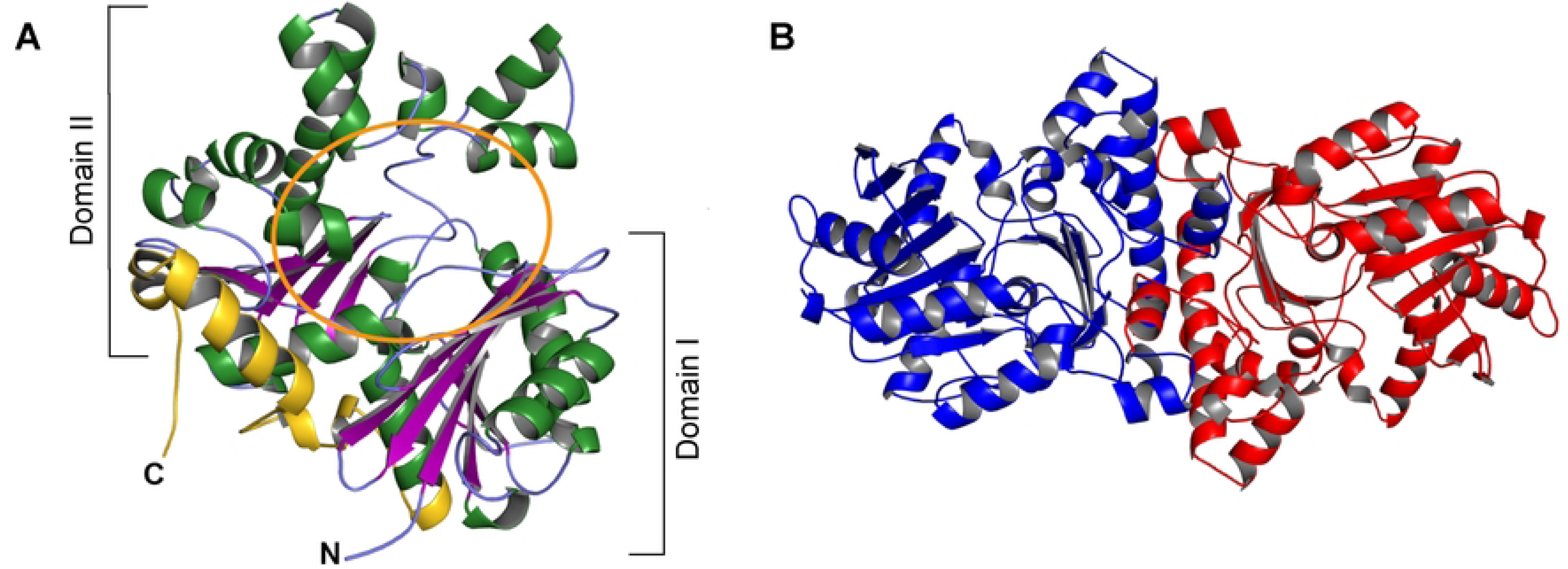
The crystal structure of TV0924. A cartoon representation of the model is shown, with helices in green, β-strands in purple, and regions without regular secondary structure in light blue. The amino-and carboxyl-termini are labeled “N” and “C”, respectively. The region encompassing putative active-site residues is circled in orange. (B) The likely dimer of TV0924. Two monomers related by a crystallographic two-fold axis are shown; one is colored red, the other blue.

Compared to other acetate kinases, TV0924 has a C-terminal extension of almost 50 amino-acids (Fig. 4, residues 401-449). In the structure, this extension comprises three helical elements, the largest of which packs on the surface of the protein well separated from the active site. The functional implications of this extension are unknown, but, given the orientation of the extension, it is unlikely to directly interact with substrates or products of the enzyme. Notably, TP0476 has the same extension, and the sequence identity between the two proteins in this region is 72% (Fig. 2).

All of our solution studies (Figs. 3B-3D) provided evidence that TV0924 was a dimer. Yet, we observed only a monomer in the asymmetric unit of the crystals. We therefore submitted the structural model to PISA (41) for the examination of molecular interfaces in the crystal. The protocol identified a single stable interface (across a twofold crystallographic axis) that featured a total of 6,110 Å^2^ in buried surface area. The model of this dimer (Fig. 4B) was subjected to hydrodynamic simulation using HullRad (42), yielding a theoretical sedimentation coefficient of 5.5 S, which is very close to the value reported above (5.387 ± 0.014 S). Given the large amount of buried surface area and the close correspondence to solution observations, this crystallographic dimer is very likely the same that was characterized in solution.

To gain information regarding the enzyme’s mechanism, we co-crystallized TV0924 in the presence of its substrate ADP and the cofactor Mg^2+^. This mixture yielded crystals under the same conditions as the apo-protein, and we acquired X-ray diffraction data from a crystal to a *d*_min_ spacing of 1.3 Å (Table 1). The overall structure of TV0924 in the presence of MgADP was very similar to that of Apo-TV0924: over the 453 comparable C_α_ atoms, there was an r.m.s.d. of 0.3 Å. Difference electron-density maps displayed evidence of an adenine-based nucleotide binding in the active site, but no density corresponding to the β-phosphate of the nucleotide was evident (Fig. 5). Also, there was apparently no electron density for an associated Mg(II) ion. The reason for the absence β-phosphate and metal ion remains unknown; it is possible that ADP was hydrolyzed during incubation with the protein in the crystallization solution or that the β-phosphate was disordered and therefore invisible in our electron-density maps. The nucleotide was modeled as AMP, and we refer to this structure below as TV0924‧AMP.

**Figure 5.**
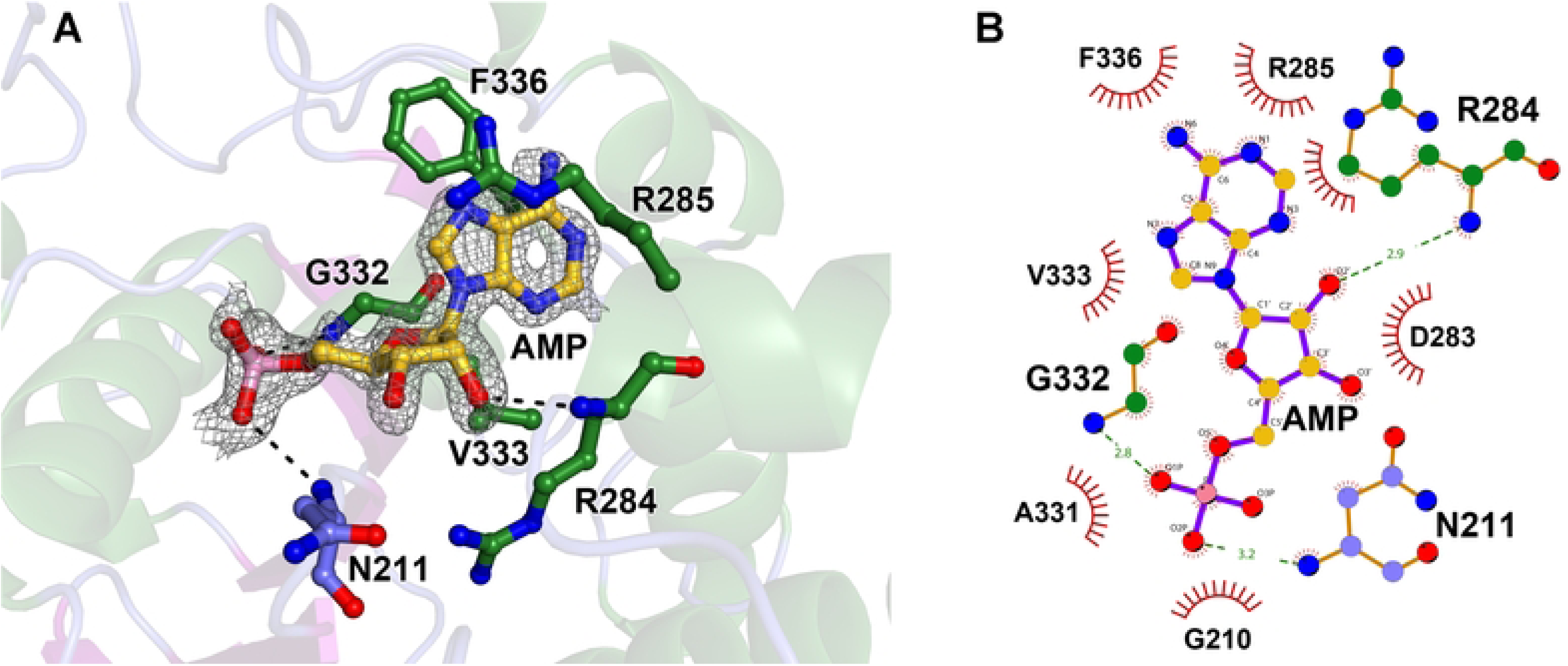
Structural characterization of AMP binding to TV0924. (A) Structural model and evidence of AMP binding. The final coordinates of the AMP and protein-nucleotide contacts. The modeled AMP nucleotide (gold carbon atoms) is shown along with relevant nearby amino-acid residues (carbon atoms colored according to secondary-structure; see legend to Fig. 4). Other atom colors are: red, oxygen; blue, nitrogen; and pink, phosphorus. Black dashes denote likely hydrogen bonds between protein and nucleotide atoms. Secondary structure of TV0924 is shown semi-transparently for clarity. Superposed on the coordinates is an *mF*_o_ – *DF*_c_ electron-density map (mesh) contoured at the 3-*σ* level; the map was calculated after omitting AMP from the final model and performing another round of refinement, and it is restricted to be within 1.2 Å of the AMP atoms. (B) Schematic view of nucleotide binding by TV0924. The color scheme is the same as in part (A), except hypothesized hydrogen bonds are shown in green, and the distances between their respective atoms are shown in Ångströms. Red spiked arcs represent hydrophobic contacts between the nucleotide and the named amino-acid residue.

The AMP molecule engaged in only a few contacts with the protein (Fig. 5). The adenine base moiety was located in a U-shaped “cradle” lined by the hydrophobic side chains of R285, V333, and F336. The base also stacked on the peptide bond between G332 and V333 (not shown). The only apparent hydrogen bonds between the nucleotide and the protein occurred either at the ribose or phosphate moieties, and these contacts were made by the protein’s main-chain atoms (Fig. 5). The electron density for the AMP overlapped with density that was similar to that observed in the apo structure (Fig. S2), and it therefore appeared that AMP was unable to displace fully the unidentified molecule. As a consequence, the occupancy of the AMP was refined, finally arriving at a value of 0.83.

### Structural comparisons to other enzymes

After searching structural databases with the model coordinates of Apo-TV0924 using secondary-structure matching (43) and distance-matrix comparisons (44), the consensus best matches (Table 3) were the acetate kinases from *Thermotoga maritima* (PDB accession code 2IIR; no attendant publication) and *Porphyromonas gingivalis* (39) (PDB accession code 6IOY). The sequences of these two proteins are shown aligned with those of TV0924 and TP0476 in Fig. 2. The models of additional enzymes, particularly propionate kinases (TdcD from *Salmonella typhimurium* (40) and PduW from *Klebsiella pneumoniae*; no attendant publication), were also good matches to the TV0924 tertiary structure (Table 3). AckA from *Methanosarcina thermophila* (MsAckA) is the best-studied bacterial acetate kinase (36–38,45), and its structural model also closely hews to that of TV0924 (Table 3, Fig. 6).

**Figure 6.**
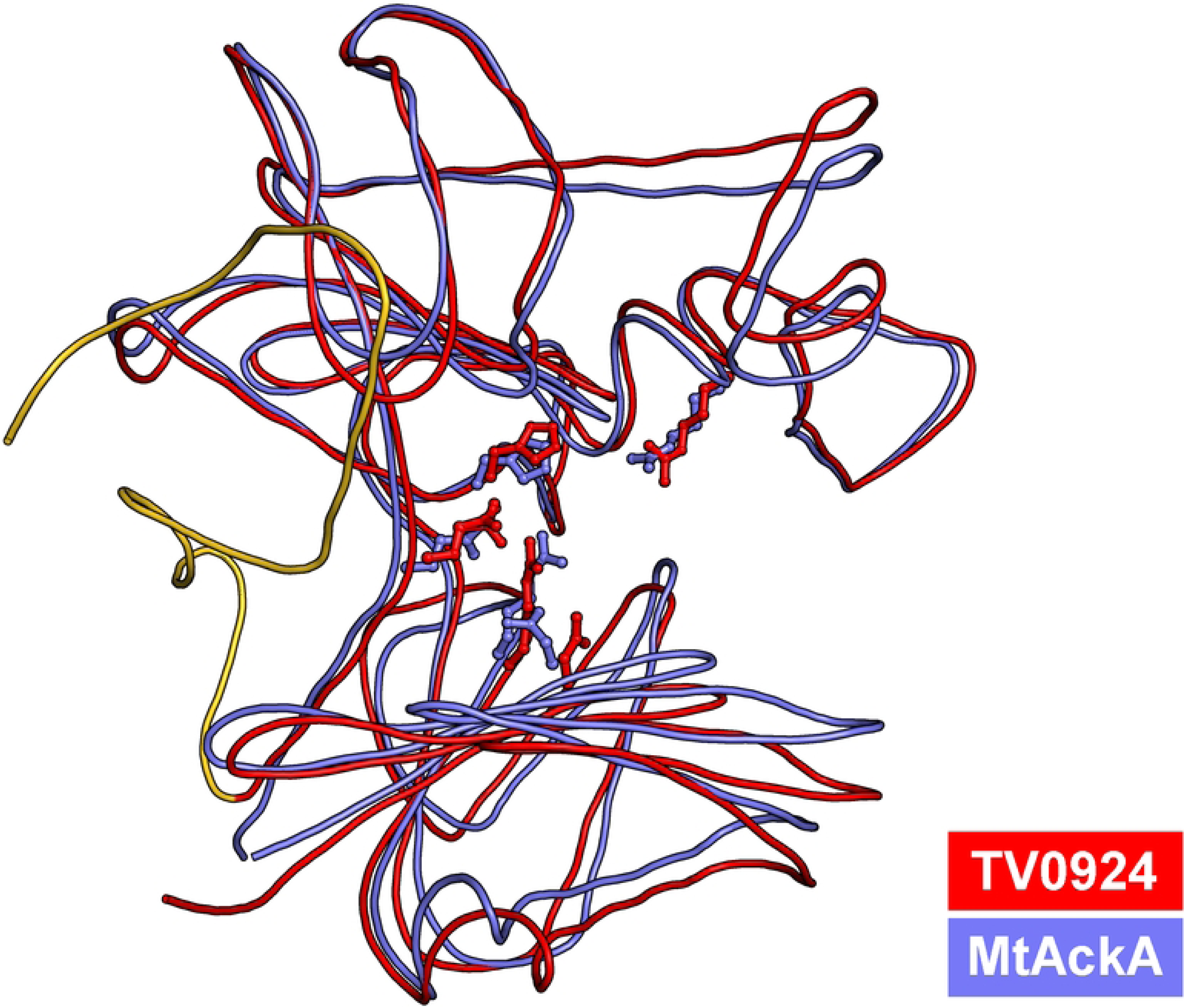
Comparison of TV0924 and MsAckA. Smoothed, cylindrical ribbons are shown depicting the main chains of TV0924 (red) and MsAckA (gray). The five putative active-site residues from the structures are shown in ball-and-stick format and are colored according to their respective structure. Residues forming the C-terminal extension in TV0924 are colored gold.

**Table 3.** Alignment statistics for selected structural models.

| Protein | SSM |  | DALI |  |
| --- | --- | --- | --- | --- |
|  | r.m.s.d. (Å) | #C<sub>α</sub> | r.m.s.d. (Å) | #C<sub>α</sub> |
| *T. maritima* AckA | 1.3 | 398 | 1.3 | 399 |
| *P. gingivalis* AckA | 1.2 | 394 | 1.3 | 398 |
| *M. thermophila* AckA | 2.0 | 390 | 2.2 | 398 |
| *S. typhimurium* TdcD | 1.2 | 382 | 1.4 | 393 |
| *K. pneumoniae* PduW | 1.2 | 388 | 1.4 | 394 |

Close scrutiny of the sequence alignment of these structurally homologous proteins (Fig. 2) reveals two striking facts. First, only the treponemal proteins harbor the C-terminal extension noted above for TV0924 and TP0476. Second, all of the important catalytic amino-acid residues identified in the *M. thermophila* protein (N7, R91, H180, R241, and E388 in the TV0924 numbering) are strictly conserved. Superposition of the *M. thermophila* coordinates on those of TV0924 shows a very close correspondence in the relative positioning of the side chains of these residues (Fig. 6).

The amino-acid residues named above feature prominently in the hypothetical catalytic mechanism for acetate kinase (Fig. 7), which was first proposed for the *M. thermophila* enzyme (38). N7 and E388 coordinate a Mg(II) ion, which in turn contacts the α-and β-phosphates of ATP. H180 is thought to be important for binding and orienting acetyl phosphate, and R91 and R241 appear to be well positioned to assist in this function. H180 and R241 (in second roles) putatively stabilize the trigonal bipyramidal transition state that is presumably formed when an oxygen atom from the ADP attacks the phosphorus atom of ADP.

**Figure 7.**
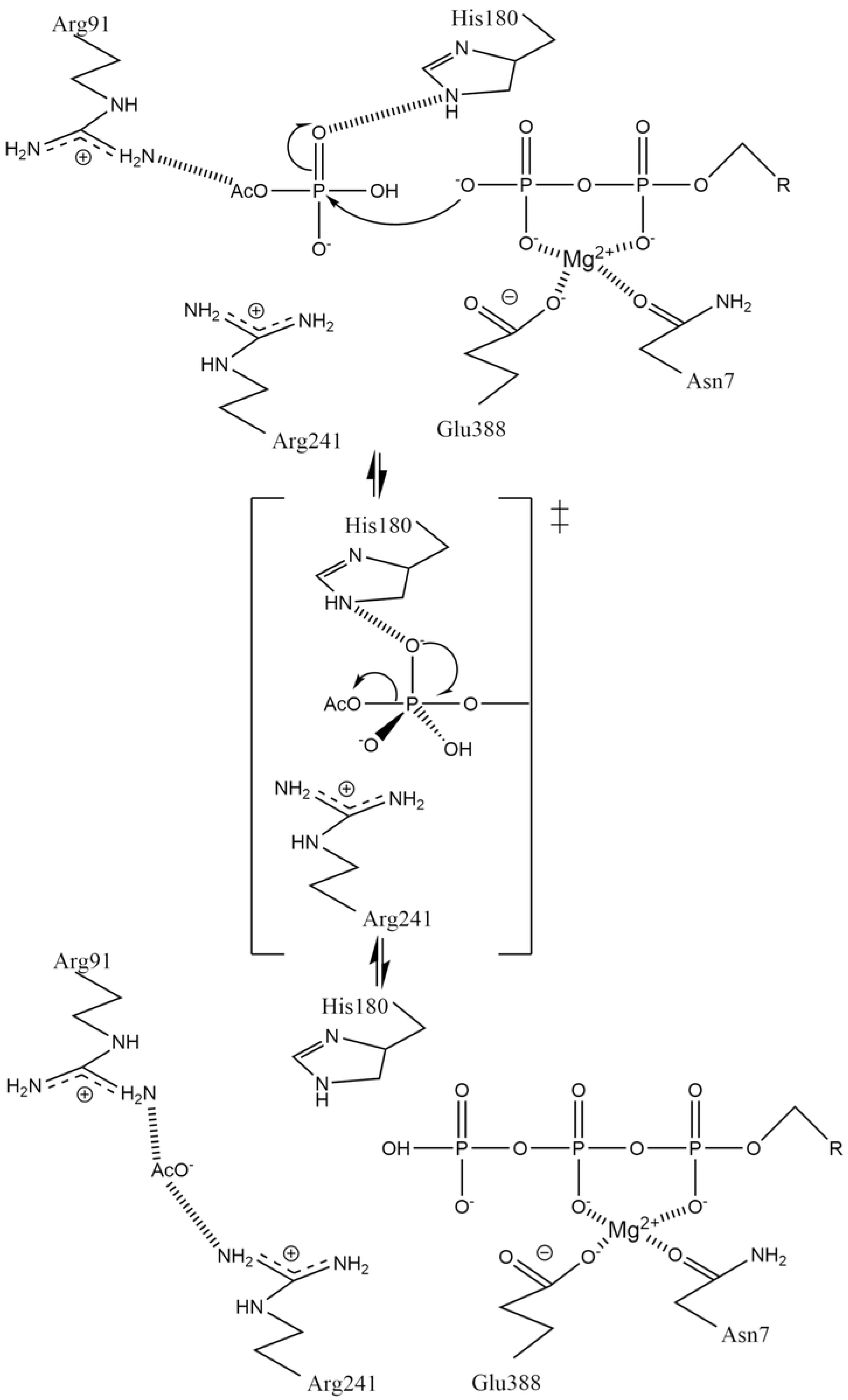
The hypothesized mechanism of bacterial acetate kinases. The assumed transition state is marked with a double-dagger and is shown in abbreviated form. R is substituted for the ribose and adenine moieties of the nucleotides. This figure was adapted from the original (Fig. 8 of (38)) under a CC-BY license (see https://creativecommons.org/licenses/by/4.0/).

### Enzyme activities of TV0924 mutants

To examine the essentiality of the above-named amino-acid residues for ATP generation by TV0924, we singly mutated them to alanine and determined the enzyme activities of the resulting proteins (Table 4). To ensure that the effects we observed were due solely to enzyme activities and not to protein stability differences, we performed stability measurements using CD, and the mutant proteins had similar characteristics to the wild-type protein (Fig. S3, Table S1). All of the mutations had strong effects on *k*_cat_, whereas the alterations regarding the *K*_M_ values were mixed. Because two substrates are utilized, henceforth we use the notation *K*_M,ADP_ and *K*_M,AcP_ to denote the respective *K*_M_ values. Compared to the wild-type enzyme, N7A and E388A had only modest effects on either of these values; R91A, and R241A featured notable increases in *K*_M,AcP_, consistent with their assumed roles of acetyl-phosphate binding and positioning. H180 appears to be a particularly pivotal residue; its mutation to alanine affected *k*_cat_, *K*_M,ADP_, and *K*_M,AcP_. That N7A did not markedly raise *K*_M_ values but strongly affected *k*_cat_ suggests that the native N7 must be critical for catalysis while its effects on substrate binding are obscure. Given its position in the active site, it seems likely that residue is needed for the ADP to adopt the optimal conformation for efficient catalysis.

**Table 4.**
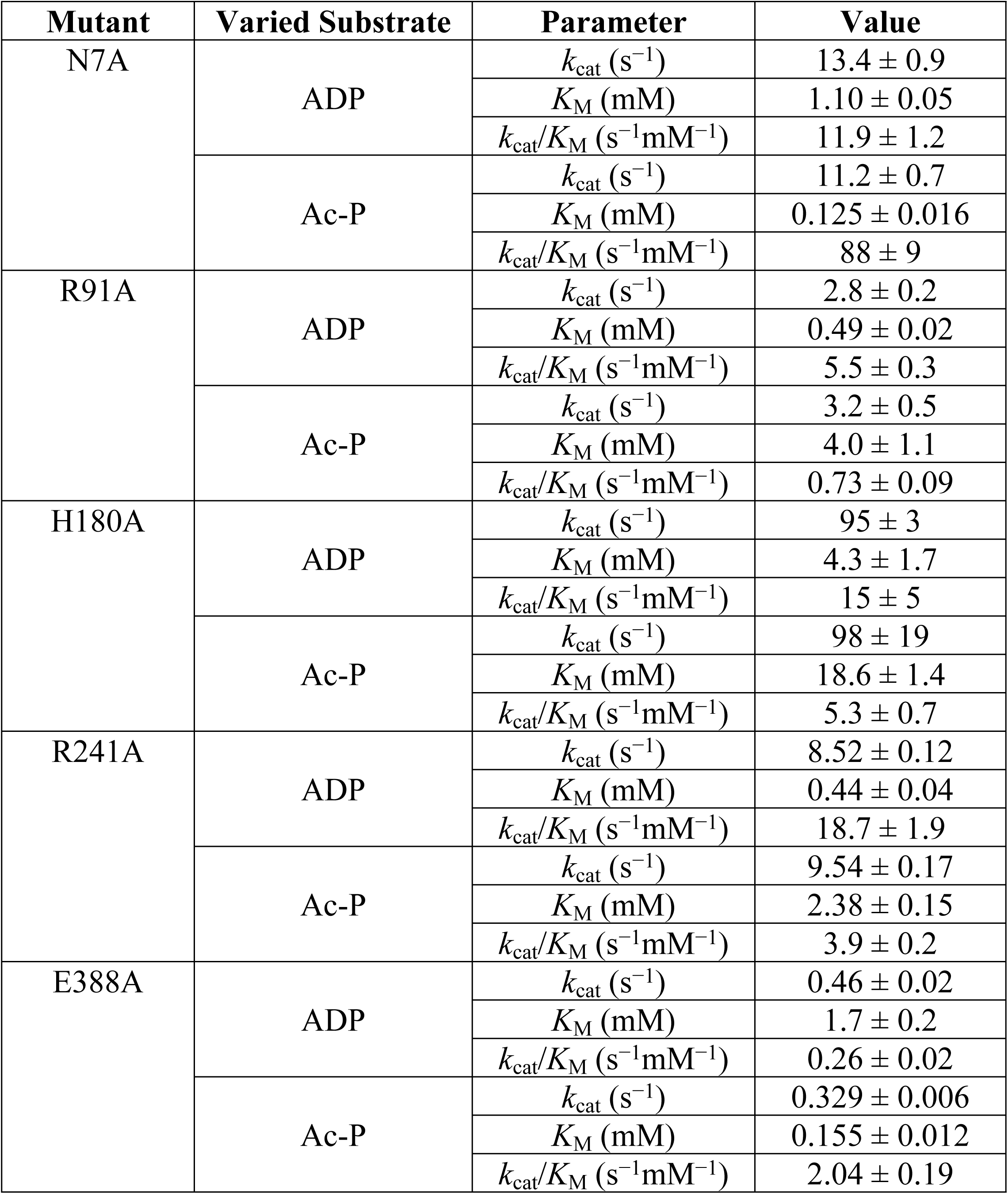
Enzymatic kinetic parameters for mutants of TV0924.

## Discussion

Our structural (Figs. 2, 4-6, Tables 1 & 3) and biophysical (Fig. 3) characterizations of TV0924 revealed a dimeric protein with a fold similar to other known acetate kinases (and, more broadly, to other sugar kinases). Enzyme assays (Table 2) demonstrated that the protein has the ability to catalyze the substrate-level phosphorylation of ADP, forming ATP using acetyl-phosphate as the phosphate donor. Given that the protein binds adenosine-based nucleotides (Fig. 5) near to amino-acid residues that are analogous to catalytic residues in other AckA enzymes (Fig. 6), and that mutation of these residues in TV0924 had significantly deleterious effects on its activity (Table 4), we concluded that the protein is a *bona fide* acetate kinase. Henceforth, we suggest that the enzyme be called TvAckA.

TP0476, the TV0924 homolog from *T. pallidum*, has a high sequence identity to the *T. vincentii* protein (Fig. 2) and also catalyzed ATP formation (Table 2). Although its solution behavior was more ambiguous (Fig. 3), TP0476 appears to favor a dimeric assembly, but it could not be crystallized to examine its overall fold. Nonetheless, its demonstrated activity and high sequence identity to TV0924 strongly suggest that the protein also carries out the functions of an acetate kinase *in vivo*, and thus we term it TpAckA.

Compared to other structurally characterized acetate kinases, TvAckA and TpAckA contain C-terminal extensions (Figs. 2, 4, & 6). In the structures of TvAckA presented herein, the extension is well ordered, forming a mostly helical arrangement that packs against the periphery of the protein well away from the active site (Fig. 4). The functional consequences of this arrangement are unknown; this adaptation was found mostly in spirochetal species, but it could be found in other (e.g. *Thermacetogenium phaeum* among many others).

With the confirmation that TpAckA is an acetate kinase, three of the four enzymes in the hypothesized acetogenic energy-conservation pathway in *T. pallidum* have been confirmed to have their putative functions. Specifically, we have investigated the D-lactate dehydrogenase (16), the phosphotransacetylase (17), and the acetate kinase (this work) in this pathway, which apparently catabolizes D-lactate, culminating in the substrate-level phosphorylation of ADP to ATP and the concomitant generation of acetate. The only remaining uncharacterized enzyme in the pathway is the pyruvate:flavodoxin oxidoreductase (PFOR), which hypothetically catalyzes the oxidative decarboxylation of pyruvate to form acetyl-CoA (Fig. 1). We have attempted to overexpress treponemal PFORs in *E. coli*, but these efforts did not yield soluble, active enzymes. Despite this shortcoming, our successful characterizations of the other three enzymes in the pathway provide strong evidence of its existence in *T. pallidum*.

## Acknowledgments

The authors thank the Structural Biology Laboratory at UT Southwestern Medical Center for support with X-ray crystallographic studies. The contents of this publication are solely the responsibility of the authors and do not necessarily represent the official views of NIAID or NIH.

## Supporting Information Captions

**Figure S1. Mass photometry of TV0924 after 63 h.** Mass photometry histograms shown at TV0924 concentrations of (A) 10 nM, (B) 20 nM, and (C) 30 nM. Blue histograms represent the number of contrast events per binned mass. Black lines are gaussian fits to the respective histogram peaks.

**Figure S2. Electron density in the active site of TV0924.** Shown are the refined positions of AMP and surrounding residues. Two electron-density maps are shown. The *gray map* shows an *mF*_o_ – *DF*_c_ map contoured at the 3-*σ* level of the omit map described in Fig. 5, but without restricting the map to be close to the depicted atoms. The *magenta map* shows the same type of density contoured at the same level for the Apo-TV0924 structure.

**Figure S3. CD-based melting curves for mutants of TV0924.** Markers are the CD data monitored at 219 nm. Lines are fits to those data using (Eq. 5). See inset legend for colors.

**Table S1. *T_m,app_* values for TV0924 mutants.** All values are presented as the fitted *T_m,app_* ± the 68.3% confidence interval.

